# Testing a mate’s brain: Courtship signals may reveal signaler decision-making capacity

**DOI:** 10.64898/2026.08.19.745821

**Authors:** Hudson Kern Reeve, Joseph Fetcho, Minhua Yan

**Affiliations:** Department of Neurobiology and Behavior, Cornell University Ithaca, New York 14853, USA; Institute For Advanced Study in Toulouse: IAST

## Abstract

It has been suggested that courtship signals reflect a potential mate’s learning ability or nervous system competence. However, there is no rigorous theory that explains how features of sexual signals represent a nervous system’s “quality”. Such a theory may provide a mechanism for mate assessment via sexual signals and offer an explanation for why courtship signals are rhythmic and stereotypic. In our paper, first we use a general model of optimal neural decision-making to show that variance in an organism’s solution time for a given fitness problem lowers the fitness gain rate; more specifically, in well-supported “competing accumulator” models of decision making, we show that noise in the slope of spike rate increase in evidence accumulators increases both reaction time and the probability of a sub-optimal decision. In conclusion, higher timing regularity leads to quicker and better decisions. This finding accords with extensive human study data showing that variance in reaction times is negatively associated with various measures of motor and cognitive performance. Thus, selection should favor individuals that require potential mates to advertise courtship signal regularity to indicate their nervous system’s general timing consistency (the timing-consistency signaling theory). The focus on signal consistency (rather than on signal duration or power) may account for why courtship signals are typically rhythmic, are often multi-modal, and why rhythmic signals are also employed in territorial contests. One of the model’s several predictions is that individuals should favor potential mates with lower noise in courtship signal features such as inter-pulse intervals.

## Introduction

Animals rely on their nervous systems to solve problems that impact their ability to survive and reproduce. They must, for example, find and compete for food and shelter/nest sites, avoid predation, and compete for mates. The ability to solve these problems depends critically upon a good nervous system. Consequently, we might suspect that animals are selected to choose mates with a high-quality nervous system so that their offspring are themselves more likely to survive and reproduce, as has been suggested by several empirical studies (Boogert NJ et al., 2011). Many have recognized that features of nervous system functionality may be represented in sexual signals and have suggested that some aspects of courtship signals (like song complexity) may provide read-outs on learning ability (Boogert NJ et al., 2008) or a more generalized aspect of nervous system competence (Boogert NJ,Fawcett TW and Lefebvre L, 2011). In other words, mating interaction is in part a neurological test where the mate-seeking individual demonstrates and the choosing individual evaluates the quality of the signaler’s nervous system. What is missing is a detailed, quantitative theory that predicts what specific measures might best represent the “quality” of the mate’s nervous system and how a mate might most efficiently advertise these specific quality attributes. Such a theory might provide a fundamental explanation for the very form of courtship signals, e.g., why they tend to be both rhythmic and stereotypic.

In this context, we searched for nervous functional measures that might effectively represent the nervous system’s quality. Human neurological studies on the relationship between simple and more generalized measures of nervous system function can be informative in this regard. Even simple measures like reaction time are related to the ability to solve a variety of complex problems (Baker LA et al., 1991;Camicioli RM et al., 2008;Luciano M et al., 2004;Luciano M et al., 2001;Metter EJ et al., 2005;Rijsdijk FV et al., 1998)

The ability to produce rhythmic movements, such as tapping, with a low interval variation also shows correlations with general measures of cognitive ability (i.e., a “good brain”) as well as with structural features of the brain (H N et al., 1996;Madison G et al., 2009;Merchant H et al., 2008;O’Boyle DJ et al., 1996;Ullén F et al., 2008). The rhythmicity is reduced in humans with neurodegenerative diseases or psychiatric disorders (Bolbecker AR et al., 2011;Da Silva FN et al., 2012;Shimoyama I et al., 1990;van den Bosch RJ et al., 1996).

Natural mating interactions have features consistent with nervous functional evaluation via simple measures. Sexual signals often involve rhythmic sound or movement production that could allow for assessment of control over interval variation. While others have recognized the potential importance of such signals in assessing nervous system function, there is no fitness-based theory that explains the mechanism underlying such a correlation and predicts what might be both strong and efficient (economical) measures of nervous system performance. Here we propose such a theory and make new, explicit predictions of the outcome of mate choice experiments in which females must choose between males differing in sexual signal timing consistency.

## Results

### The timing-consistency signaling theory

#### An abstract model

The central idea behind our model of courtship signaling is that courtship signals provide an individual with information on the decision-making ability of the potential mate’s nervous system. We show that higher timing regularity leads to (i) a lesser latency to decision time for a fitness-maximizing organism, as well as (ii) a greater probability that the organism will make the correct, i.e., fitness-maximizing, decision. Thus, a mate-seeking individual that favors potential mates with greater neural timing regularity will obtain mates that can make both quicker and better decisions, traits that can be inherited by the offspring of the mating. First, we develop an abstract optimality model of how variance in decision time will affect an individual’s rate of fitness accrual:

An organism’s behavior is built by natural selection to accrue fitness benefits by solving fitness-related problems in a time-efficient manner. Suppose that an organism confronts a problem whose solution will generate a fitness benefit. Natural selection should favor organisms that obtain such benefits as rapidly as possible because this would increase the overall rate of fitness accrual and thus the cumulative fitness gain from solving repeatedly encountered problems. However, there is a trade-off. The likelihood of finding a better solution for the current stimulus context increases with the time invested in assessing the problem and generating and comparing potential solutions. In other words, the greater the time investment, the higher the quality of the solution but the less the amount of time available for accruing fitness benefits from solving other problems. Therefore, to maximize the rate of fitness accrual, an organism must balance between the number of fitness problems solved and their gain from each problem.

We can model the optimal investment time in the following way. Let *t* be the time invested to extract a benefit *B(t)* for an encountered problem. The time *t* includes the time to assess the problem, compare potential alternative solutions, and implement the optimal solution. The benefit *B(t)* is an increasing function of the investment time *t*, but at some point we expect *B*(t) to be decelerating, i.e., *B*’’(*t*) < 0. The latter is true because increasing investment time should produce declining fitness benefits: Since the probability that a better choice than those that have already been assessed will be found must decrease as investment time increases, there must eventually be a smaller further fitness gain for investing more time.

Natural selection should act to maximize the long-run expected fitness accrual rate *B(t)*/*t*. The solution time *t* will vary within an individual due to stochasticity in the operation of the nervous system, and such stochasticity may vary across individuals in a heritable way. It follows from the elementary renewal theorem applied to repeated solution of fitness problems that selection should maximize the long-run fitness gain rate R:

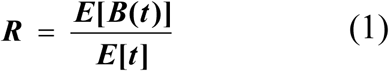

where *E[B(t)]* is the expected fitness gain from a problem solution and *E[t]* is the mean time invested in obtaining a solution.

Now let’s suppose the mean value of *t* (= *E*[*t*] = *m*) is under the control of the organism and is the target of natural selection but that the variance in *t* (= *v*) reflects an inherent, irreducible inefficiency in neural control. We are particularly interested in how the variance *v* in investment time will affect the fitness accrual rate when the organism employs the mean investment time *m* that maximizes (1).

We can use a Taylor expansion to expand the function *B(t)* in the vicinity of the mean value of *t* = *m* (ignoring terms of higher order than 2 here and below) to obtain.

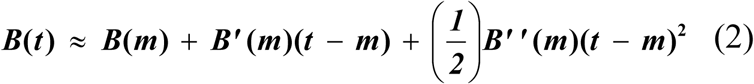

Taking the expectation of (2) over random variation in *t* yields

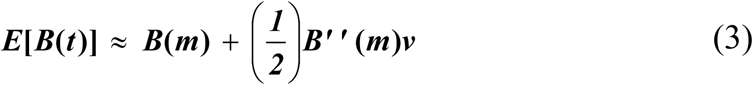

since *E*[(*t*-*m*)] = 0 and *E*[(*t*-*m*)^2^] = *v*.

Substituting (3) into the numerator of (1) and setting *E*[*t*] = *m* yields, as the approximate fitness accrual rate *R* to be maximized,

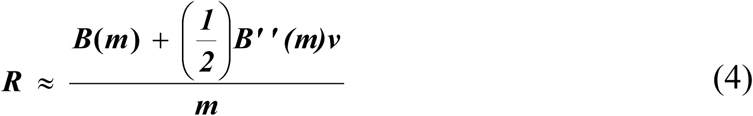

The mean investment time *m* = *m*\* that maximizes *R* must satisfy d*R*/d*m* = 0, and d^2^*R*/d*m*^2^ < 0, at *m* = *m*\*. The latter condition for a maximum entails that *B*’’(*m\**) < 0, i.e., that the optimal mean investment must correspond to a decelerating (i.e., “diminishing returns”) region of the *B*(*t*) function (Fig. 1).

**Figure 1.**
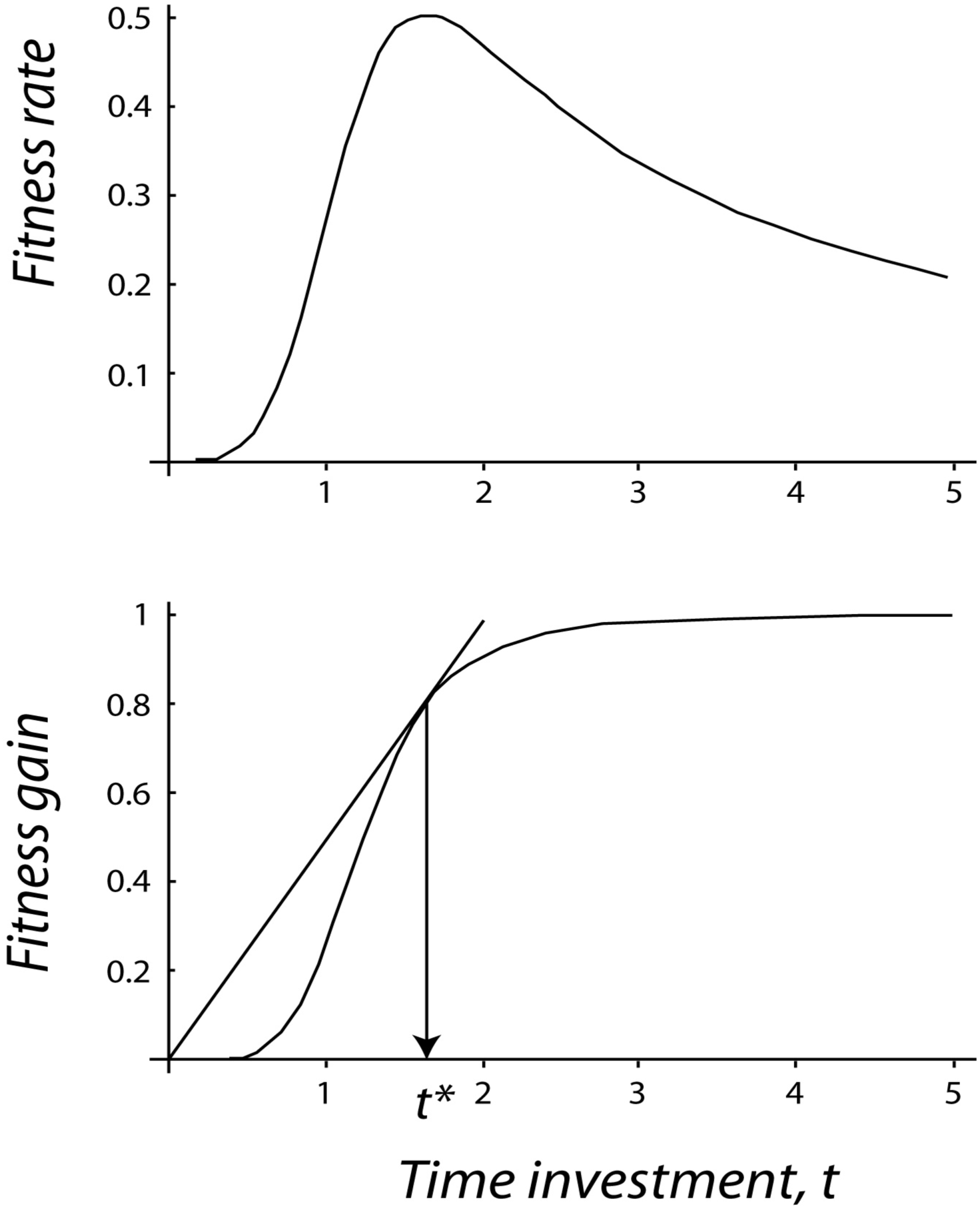
Optimal time investment in neural decision-making: effect of variable decision times. (Top) The organism’s nervous system is assumed designed to optimize the time *t* invested in reaching and implementing a decision about how to solve a problem, given that there is eventually a diminishing fitness gain to time invested and that selection maximizes the fitness gain rate. The graphical solution for finding the optimal time *t*\* is found by drawing the line that passes through the origin and just touches the fitness gain curve, then dropping a perpendicular, as shown. (Bottom) The fitness gain rate is at a peak at the optimal investment time. Thus, any variation in the decision time around the optimum will only reduce fitness.

Now what we seek is how the variance *v* will affect *R* given that the organism employs the optimal *m*\*. Mathematically, this means that we are seeking d*R*/d*v* at *m* = *m*\*. Using the approximation in (4), we have:

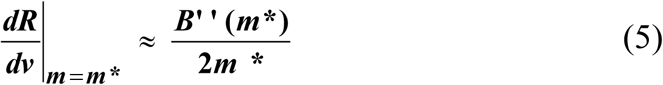

Since *B*’’(*m\**) < 0 if *m*\* is to be a maximum (see expression 5), it follows that d*R*/d*v* is negative and thus organismal fitness rate must decline as the irreducible variance *v* in investment times increases. In other words, organisms that exhibit a higher variance in decision times will experience lower mean fitness accrual rates. (Equivalently, higher coefficients of variation in decision times will lower fitness.)

Interestingly, expression 5 entails that there will be stronger selection against the variance *v* when the optimal decision time *m*\* (see Appendix A) is shorter. Therefore, our model predicts that in species in which decisions must be especially quick, there will be an especially large benefit for obtaining mates with a lower neural variance *v*.

### The effect of neural noise on reaction time and probability of a correct decision in neural models of decision making

The above is an abstract model of optimal decision making not grounded in any particular neural mechanism underlying decision processes. However, over the last decade or so, evidence from both the psychological and neurobiological literature has strongly supported what are called the drift-diffusion (DD) or closely related Linear Ballistic Accelerator (LBA) models of the underlying decision process (Ratcliff, 2008; Brown and Heathcote, 2008). We will not distinguish between DD and LBA models, which make very similar predictions (Donkin C et al., 2011), but briefly describe a simple version of what they have in common: Suppose that there are two competing neural circuits corresponding to two different behavioral decisions, A and B. In each circuit, the spike rate of an output neuron increases at some mean rate, the actual rate being subject to random variability (Brunton BW et al., 2013;Churchland AK et al., 2011), and a decision is “reached” when the actual spike rate climbs to some critical threshold. Which of the two competing decisions is enacted depends on which circuit hits its threshold first. Interestingly, these decision models have been shown to theoretically maximize reward rate, as assumed in our abstract model above (Bogacz R et al., 2006).

Now we show that in the DD /LBA multiple accumulator models, a greater variance in neural timing, in this case, variance in the slope of spike rate increase for the “winning” circuit, increases the average time until the ultimate decision is made. Let the slope of spike rate increase in this circuit be equal to *s*, which is subject to noise described by some variance *v*. If the circuit’s threshold is *T*, a decision is made at time *t* = *T*/*s*. Using a second-order Taylor expansion around the expected value of *s*, *sav*, the expected time E(*t*) is approximately equal to

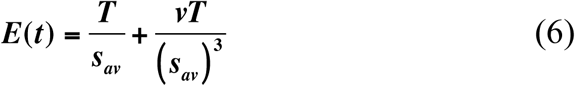

As the variance *v* in the spike rate increases, the average time to a decision increases, as in the more abstract optimality argument (Fig. 2). Thus, neural timing irregularity increases the time to make a decision.

**Figure 2.**
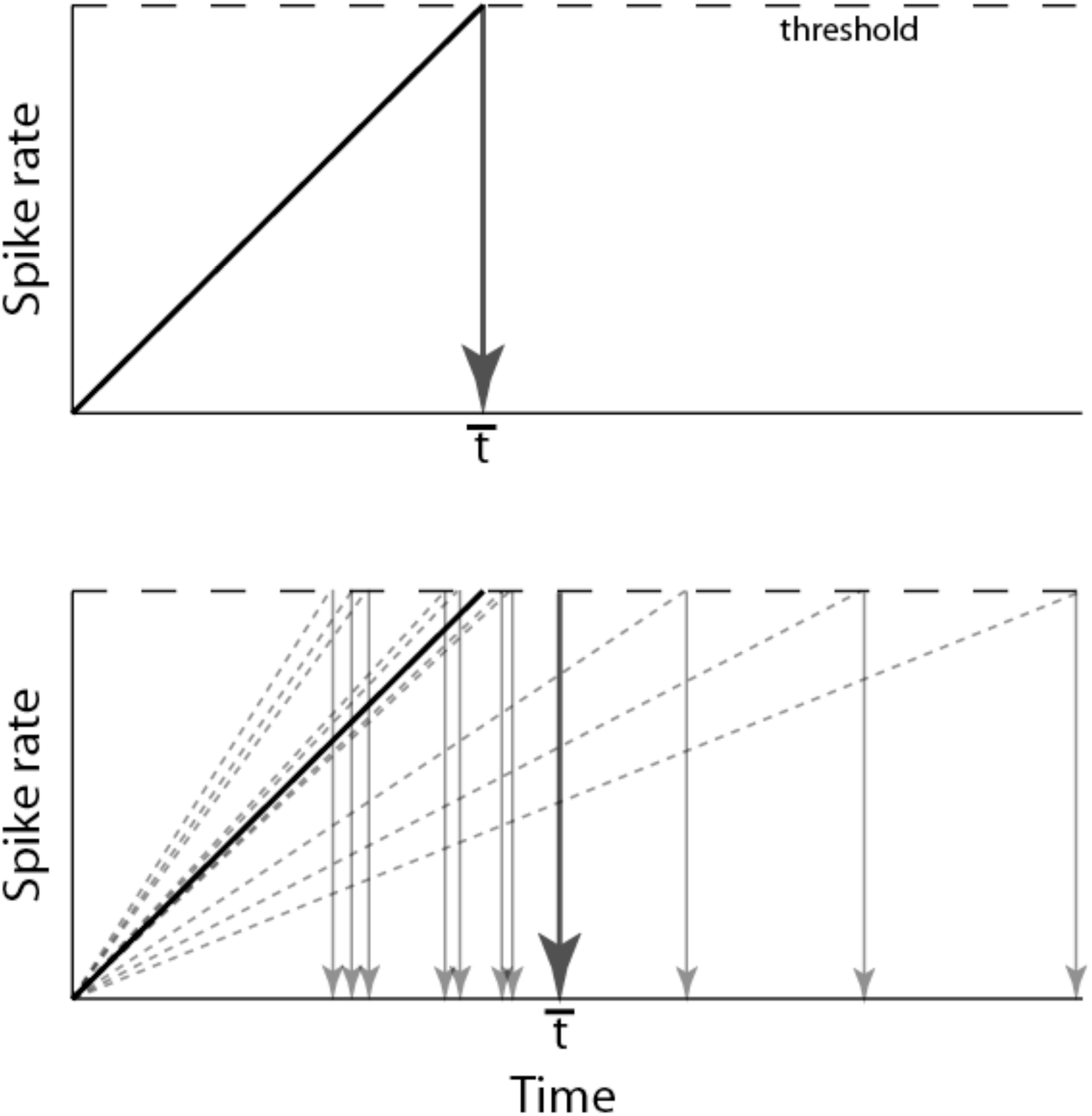
Effect of neural noise in DD/LBA models of decision making. (Top) The spike rate of an accumulator circuit output neuron rises without noise until a threshold is reached, and the decision option corresponding to the accumulator is made. (Bottom) Noise in the mean rate of spike rate increase increases the mean time until a decision is made.

**Figure 3.**
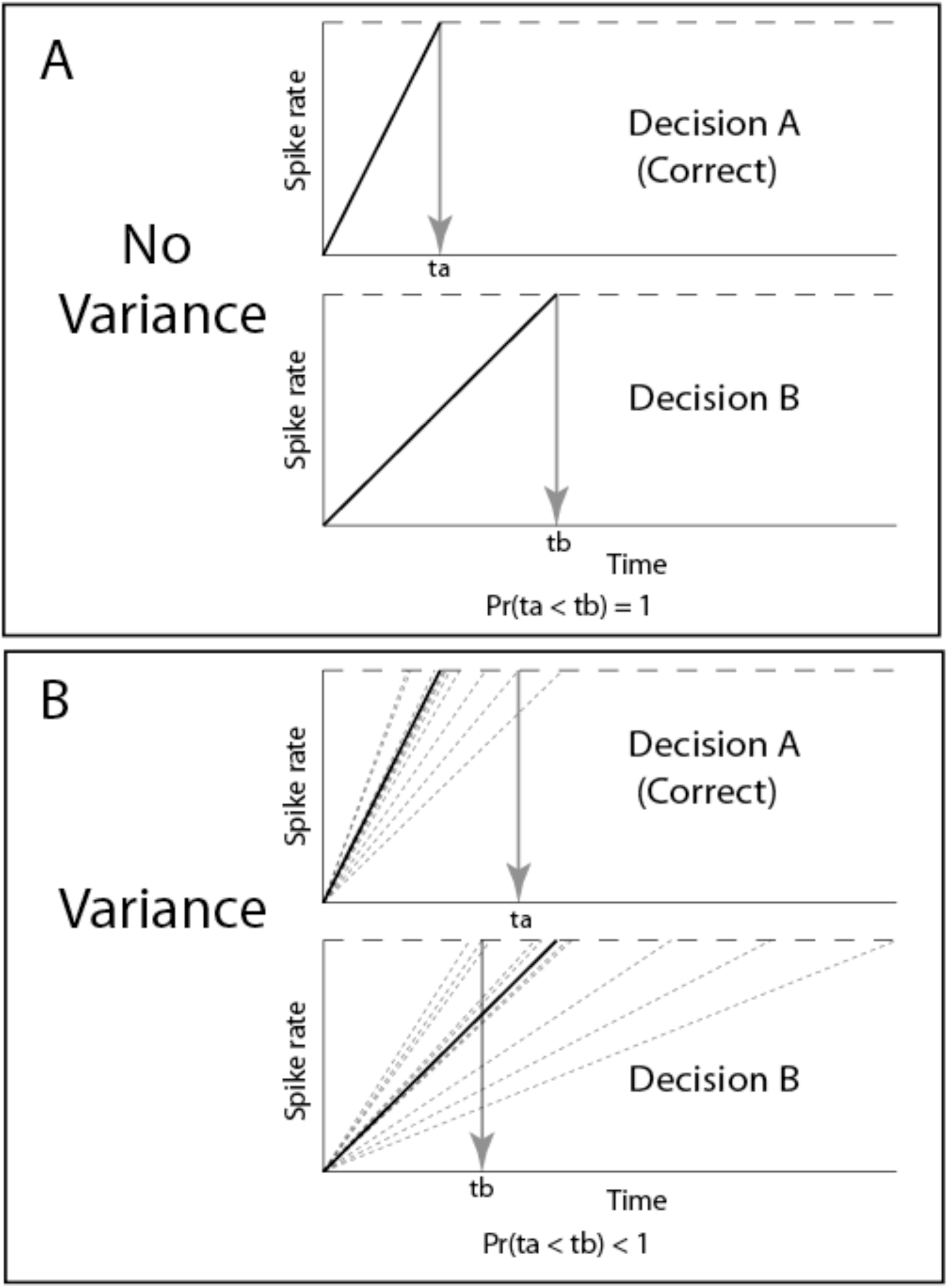
Noise in the rate of spike rate increase increases the probability that the optimal (fitness-maximizing) decision will not be made. (A) Without noise, the greater slope of the rate of spike rate for correct decision A means that the correct decision A will be made with certainty. (B). With noise in the mean rate of spike rate increase, there is now some non-zero probability that the incorrect decision B will be made first.

Next, we examine the effect of neural noise on the probability that an organism will make a correct decision in DD/LBA mechanisms. Suppose again that there are two competing neural circuits corresponding to two different behavioral decisions, A and B. The first circuit, corresponding to the correct (i.e., fitness maximizing) decision A, exhibits a slope of spike rate increase equal to *sA*. The second circuit, corresponding to the incorrect (i.e., not fitness maximizing) decision B, exhibits a slope of spike rate increase equal to *sB*. In a competing accumulator (DD or LBA) model, the probability *P* that the correct decision A is enacted is the same as the probability that the time *tA* to threshold *TA* in circuit A is less than the time *tB* to threshold *TB* in circuit B. The time to threshold in circuit A is just *tA* = *sA*/ *TA* and that in circuit *B* is just *tB* = *sB*/ *TB.* Thus, we seek the probability that *sA* > *sB* (*TA* / *TB*), or, equivalently, the probability *P* that *x* > 0, where *x* = *sA* -*sB* (*TA* / *TB*). If we make the usual assumption that *sA* and *sB* are independently normally distributed with the same variance *v*, then *x* will be normally distributed, say with mean *u* > 0 and standard deviation *ß*. (*u* > 0 follows from optimization of the nervous system by natural selection, with the result that the correct decision will be made more than 50% of the time) The standard deviation *ß* reflects noise in the slope of spike rates (because *ß^2^* = *v*[1 + ( *TA* / *TB*)^2^], where *v^1/2^* is the standard deviation of the spike rate slope).

It follows from the above that the probability of a correct decision *P* is equal to

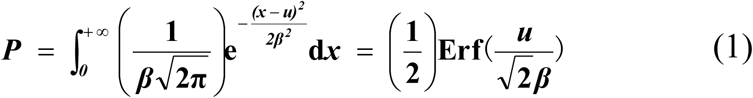

where *Erf* denotes the error function. Thus, *dP/ds* is equal to

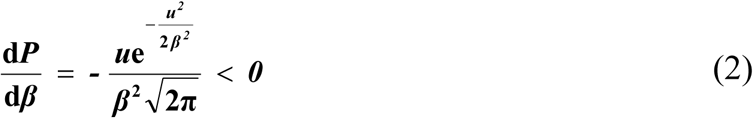

Since d*P*/d*β* < 0, it follows from the above that the probability of a correct decision will always decrease as the variance in spike rate slope increases. Thus, reduced error variance in spike rates should tend to increase fitness by leading to enhanced likelihood of correct decision making (Fig. 2). In sum, neural noise both increases mean reaction time and reduces the probability of a correct decision in DD/LBA models of decision making at the neural level. The latter result is also obtained from a more exact model of decision making involving *n* competing accumulators (Appendix B).

### How can individuals assess the timing regularity of potential mates?

Now suppose an organism is attempting to assess the overall timing regularity of a potential mate, e.g., the coefficient of variation (CV) in decision times since lower heritable values of CV are more desirable in mates, because they can make both faster and better decisions. One way that a choosing individual might do this would be to simply observe problem-solving latencies of potential mates across a large number of problems and infer CV for these prospective mates directly. However, such an assessment strategy is likely to be very time consuming, hence costly, to the choosing individual. A second possibility is that the choosing individual requires for mating that the potential mate demonstrate its CV by repeatedly solving problems in the presence of the choosy individual. However, solving such problems is likely to be quite costly in time and energy to the performing potential mate, making it less beneficial to perform if the potential performer has other options, and costly in time for the choosing individual. A further problem is that the two preceding assessment strategies may not be very reliable, as a choosing individual inferring a reaction time or problem-solving interval would have to accurately ascertain the moment when the problem-solving behavior was initiated, which may be a hidden event. However, there is a third possibility that avoids all of the above problems and it might lead directly to the evolution of rhythmic courtship signals. The choosy individual might infer CV easily directly and rapidly by measuring the CV in a relatively low-cost output of a nervous system rhythm generator such as a vocalization-producing circuit if that signal reflects overall variance in neural control mechanisms. That one output circuit might reflect at least some features of nervous system-wide timing inefficiency is plausible because different circuits share structural and functional elements used to build neurons and networks, such as ion channels, myelin, and synaptic proteins. In addition, the timing regularity of different neuronal subsystems might also be sampled directly by assessing variation in multimodal displays that engage different brain regions. In sum, to assess nervous system timing regularity of a prospective mate, a choosy female need only monitor the variability in the times between repeatedly produced, stereotypical, signal components such as pulses or notes in a song.

If nervous system timing is heritable, the above arguments therefore imply that females (for example) should favor as mates males that produce highly regular rhythmic signals, i.e., signals with a low index of dispersion. For example, if courtship signals consist of acoustically generated pulses, selection should favor evenly-spaced pulses, i.e., inter-pulse intervals exhibiting low CV’s. Such low variation may serve as an indicator not only of the competence of the underlying rhythm generator but of overall nervous system performance ability (encoded by CV). Low nervous system CV might be signaled in a variety of kinds of courtship signals, such as by regularity in repeated visual displays or even in highly consistent song-type repetition, since reproducibility of song-types must depend on constancy of timing relationships among multiple neural inputs (de Kort et al. date?).

If a female is to assess the quality of a potential mate by examining variation in a signal, then the nervous system of the female must itself be good enough to assess that variation. By what neural mechanism might females assess a signal CV from the rhythmic courtship signals of potential mates? One possibility is opened by the discovery of so-called “counting” neurons in the anuran midbrain (Edwards CJ et al., 2002). These neurons respond only after the presentation of a particular number of rhythmic pulses with interval variation below some fixed value. One long or short interval (even as little as 2 msec off the best interval) in a pulse sequence resets the counting (Edwards CJ,Alder TB and Rose GJ, 2002;Edwards CJ et al., 2008;Leary CJ et al., 2008). This ability to monitor consistency in pulse intervals at the single cell level could provide a neural substrate for assessing signal CV.

Since for this to work, good brains matter on both sides, (the females need to be able to reliably assess CV, as much as the males must produce low CV signals) a female’s choice of a competent male brain may therefore yield a self-reinforcing process: Females with brains that are able to make the best distinctions in timing measures from different males will mate with the “best-brained” males with the consequence that the offspring receive high quality genes for brains from both parents (in the parlance of sexual selection theory, both good gene effects and “Fisherian” effects can power the co-evolution of the female preference and the male signal). A particularly intriguing possibility is that a single gene that increases nervous system timing efficiency may increase both the rhythmicity of a male’s signal and the ability of a female to assess the rhythmicity of the signal. These pleiotropic effects of a single gene generate a powerful genetic coupling between a female preference and a preferred male trait that will propel both toward fixation in the population much more rapidly than will conventional Fisherian selection acting on separate, unlinked, genes.

## Discussion

### Implications of the model

The neural competence model of courtship signaling has a number of intriguing implications, which we here delineate:

1. The model predicts that organisms confronting fitness problems with typically short solution times (low *m\**) will be under especially intense selection to produce signals with low CV, as the fitness disadvantage of temporal variance in the neural system increases as the mean solution time *m*\* decreases (see appendix A). A female assessing a male’s neural competence should demand greater male investment in the signaling of CV as CV becomes more important in determining fitness. Thus, species typically confronting fitness problems requiring rapid solution might be expected to have the most elaborate rhythmic displays or songs designed to convey low CV’s. Investment in increasing the regularity of courtship signals should be especially high when organisms must continually and rapidly adjust motor behavior to survive.
2. The existence of sexual signaling through multiple sensory modalities is highly compatible with the timing consistency signaling theory. The theory predicts that sexual signaling will often occur in multiple modalities if CV is most reliably assessed when signaled through multiple modalities. Multiple cues are now known to interact synergistically to increase perceptual sensitivity through neural integration of parallel sensory pathways (e.g., Gu et al. 2008). In addition, multi-modal signaling may serve to signal the indices of dispersion associated with different major parts of the nervous system, such as timing elements in the spinal cord (e.g., for visual or vibrational displays involving limbs) and timing elements in the brain (for acoustic displays).
3. The timing consistency signaling theory presented above assumes that the signaling interaction is between potential mates. It is natural to ask what the theory might predict for strictly competitive signaling interactions such as communication between rival males at a territorial boundary. In this case, signals that advertise overall nervous system competence are important as well because they are likely to reflect the ability of an opponent to react quickly and in the appropriate ways to win a fight. Rival males might use the displays to assess their chances of winning and avoid a costly actual direct conflict. Thus, we might expect a signal of a competent brain by one party to be immediately followed by similar signals by the other party. Indeed, this idea can account for why song-matching appears much more common in avian male territorial songs than in their courtship songs (Bradbury and Vehrencamp).

### Evidence for the timing consistency signaling theory

We next turn to current evidence for the major assumptions and predictions of the neural-competence signaling model:

1. A critical prediction of the neural-competence signaling model is that both reaction speed and stereotypy (low CV in nervous system timing) should positively correlate with overall nervous system function, e.g., overall problem-solving ability. This connection has been most thoroughly explored in humans. Studies have repeatedly demonstrated that performance in tests of cognitive ability in humans is negatively correlated with mean reaction time in a number of simple motor and discrimination tasks (Brody, 1992; Deary et al. 2001). Moreover, the positive association of cognitive performance with survival in humans appears to be associated with the IQ-reaction time correlation (Deary IJ and Der G, 2005). In the context of the abstract decision model, slowed reaction time might be the consequence of a more-slowly increasing gain function, *B*(*t*) for some individuals (see Fig. 1). However, another strong possibility is that individuals vary in their neural noisiness, which should lead to increased reaction time for individuals with greater intrinsic neural noisiness (equation [6]).

Most importantly, our decision model assumes that intra-individual CV in reaction latency is variable across individuals and predicts that lower CV should be associated with greater nervous system fitness-accrual efficiency. Intriguingly, the variance in reaction times within an individual also is significantly negatively correlated with individual performance on cognitive ability tests, often more strongly than is mean reaction time (Brody N, 1992;Hultsch DF et al., 2000;Larson GE and Alderton DL, 1990), in accordance with our decision model.

Even more intriguingly, individuals who are able to tap their fingers with a higher degree of stereotypy also tend to have better performance in tests of cognitive ability, even when the reaction times are on the millisecond level (Ullén F,Forsman L,Blom Ö,Karabanov A and Madison G, 2008). The variation in tapping stereotypy negatively correlates with prefrontal white matter volume (Ullén F,Forsman L,Blom Ö,Karabanov A and Madison G, 2008). Psychiatric diseases such as schizophrenia, depression and borderline personality disorder as well as neurodegenerative diseases such as Alzeheimer’s disease are associated with larger intra-individual variability in reaction times (Gorus E et al., 2008;Kaiser S et al., 2008;Ullén F,Forsman L,Blom Ö,Karabanov A and Madison G, 2008). As expected, tests for neural deficits in clinical neurological examinations often focus on the ability to produce regularly rhythmic outputs. These results establish the kind of empirical connection between ability to exhibit rhythm stereotypy and cognitive performance that is postulated by the neural-competence signaling hypothesis. In sum, the negative correlation between cognitive performance and reaction times and the positive correlation between finger-tapping regularity and cognitive performance in humans are simply explained by our model: High neural timing regularity leads both to quicker reaction times and to better choices in decision problems.

Our model assumes that the connections between reaction times, rhythm stereotypy, and behavioral performance are heritable. Indeed, both reaction times and the correlations between reaction times and intelligence test performance have been found in human twin studies to be highly heritable (Baker LA,Vernon PA and Ho H-Z, 1991;Rijsdijk FV,Vernon PA and Boomsma DI, 1998) . One mechanistic contributor to such variability might be axonal conduction velocity, which shows heritable variation in human and mice (Reed TE, 1988;Rijsdijk FV and Boomsma DI, 1997;Rijsdijk FV et al., 1995;Vernon PA and Mori M, 1992). Changes in ion channel properties, densities and myelination could alter timing widely throughout the nervous system to alter mean and variance in processing speed for many tasks. Properties such as white matter integrity also show heritable variation that might in part reflect underlying genetic variation in myelination that could influence mean and variance in processing speed (Chiang M-C et al., 2009).

The strongest direct tests of our neural competence signaling theory would involve the following pieces of evidence from a study species: (1) Courtship signals that are less variable (i.e., exhibit lower CV) in some measure should be more attractive to the opposite sex. For example, there is already some evidence that song consistency is favored by sexual selection in the context of either mating or fighting (de Kort SR et al., 2009), as predicted by the model if song consistency reflects neural timing regularity. In contrast, Gerhardt and Watson (1995) (Gerhardt HC and Watson GF, 1995) reported that female grey tree frogs did not discriminate between male songs of low and high temporal variability. However, they did show that females did significantly prefer males with lower song variability in one experiment and females exhibited such preferences in four of the five other experiments in which the preferences were not statistically significant, possibly due to small sample sizes. Thus, it’s unclear whether the latter data falsifies timing-consistency signaling theory. We predict that female preferences for lower variability courtship displays will be found for many animal taxa exhibiting rhythmic courtship displays, even after controlling for display features relating to signal power and signaler body conditions. (2) Males that produce less variable signals (e.g., male crickets with lower CV’s in inter-pulse intervals) should ultimately be more efficient at solving multiple other motor problems relating to survival and reproduction, such as ability to catch prey or flee predators. (3) The offspring of the above males should also excel at the same tasks and exhibit heritability in CV. If the theory is supported, it may not only provide a general explanation for the rhythmic form of courtship signals, but also open the door for finding deep connections between the detailed design of courtship signals and the most critical behavioral problems confronting members of the species exhibiting those signals.

## Author contributions

H. Reeve originated the underlying hypothesis and developed the abstract decision model. M. Yan refined the model as applied to drift-diffusion neural processes. J. Fetcho originated the idea that this could explain the results of IQ-reaction time and IQ-related finger-tapping studies.

## Appendix A

### Proof that selection against neural noise becomes stronger for shorter optimal decision times

According to expression (5), the strength of selection against neural noise *v* is equal to 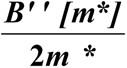 , where *m*\* is the optimal decision time and *B*[*m*\*] is the function that gives the fitness payoff for the optimal decision time.

Differentiating with respect to *m*\*, we obtain

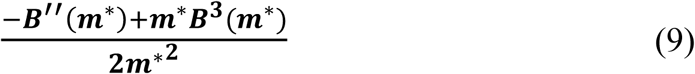

For small *m*\*, the expression (9) will tend to be positive because *B’’(m\**) is less than zero and the *m*\**B*^3^(*m\**) term can be neglected. Thus, as *m*\* gets smaller, expression (5) will become of larger magnitude and consequently, selection against the variance, *v*, will become stronger.

## Appendix B

### Proof that the probability of a correct decision decreases as neural noise increases in a drift-diffusion model with *n* options

Usher et. al., (2002) computed the probability that the correct choice out of *n* alternative options will be chosen in a race among *n* competing accumulators, where the activation of the “correct” accumulator at increases a rate µ (called the drift parameter), but is subject to Gaussian noise with mean 0 and variance σ^2^ . Under the assumption that only the drift parameter of the correct accumulator is greater than zero, with those of the competing accumulators being set to zero by natural selection, the probability of correct choice is approximately equal to

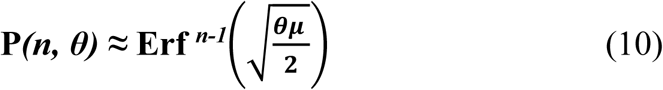

where θ is the threshold, and µ is the drift parameter (Usher et al. 2002). These authors scaled θ and µ in equation (10) in units of σ, making them dimensionless. To put σ back in the probability of a correct choice, let θ’ be the actual threshold in activation units (e.g., the spike rate of the accumulator output neuron). Then θ’ is equal to θσ, so that θ = θ’/σ . Likewise, the drift parameter in units of actual activation, µ’, is equal to µσ, yielding µ = µ’/σ. Substituting these values in for θ and µ, in equation (9), yields

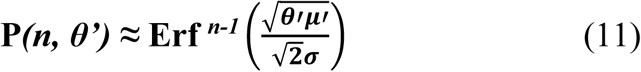

The right hand-side of (10) must increase as σ declines since the error function always increases as its argument increases. Therefore, it is readily seen that the probability of a correct choice will decline as σ increases.

